# Automated cell detection for immediate early gene-expressing neurons using inhomogeneous background subtraction in fluorescent images

**DOI:** 10.1101/2024.11.07.622525

**Authors:** Hisayuki Osanai, Miari Arai, Takashi Kitamura, Sachie K. Ogawa

## Abstract

Although many methods for automated fluorescent-labeled cell detection have been proposed, not all of them assume a highly inhomogeneous background arising from complex biological structures. Here, we propose an automated cell detection algorithm that accounts for and subtracts the inhomogeneous background by avoiding high-intensity pixels in the blur filtering calculation. Cells were detected by intensity thresholding in the background-subtracted image, and the algorithm’s performance was tested on NeuN- and c-Fos-stained images in the mouse prefrontal cortex and hippocampal dentate gyrus. In addition, applications in c-Fos positive cell counting and the quantification for the expression level in double-labeled cells were demonstrated. Our method of automated detection after background assumption (ADABA) offers the advantage of high-throughput and unbiased analysis in regions with complex biological structures that produce inhomogeneous background.

**Highlights:**

- We proposed a method to assume and subtract inhomogeneous background pattern. **(79/85)**
- Cells were automatically detected in the background-subtracted image. **(71/85)**
- The automated detection results corresponded with the manual detection. **(73/85)**
- Detection of IEG positive cells and overlapping with neural marker were demonstrated. **(85/85)**

## 1. Introduction

Immediate-early genes (IEGs) are rapidly and transiently transcribed in response to extracellular signals, and have been shown to be involved in long-term synaptic plasticity and memory (Ma et al 2023, Sheng & Greenberg 1990). c-Fos is one of the first characterized IEGs (Morgan et al 1987, Sagar et al 1988), and is critical for long-term-potentiation and consolidation of long-term memory (Fleischmann et al 2003, Katche et al 2010). Since IEG expression reflects synaptic plasticity induced by neuronal activity (Tonegawa et al 2015), IEGs have been widely used as endogenous markers of neuronal activity (Hoffman et al 1993, Minatohara et al 2015, Yokose et al 2024, Yokose et al 2023) allowing the tracking of memory engram cells (Barth et al 2004, Denny et al 2014, Kitamura et al 2017, Kitamura et al 2015, Liu et al 2012, Marks et al 2022, Reijmers et al 2007, Tanaka et al 2018, Terranova et al 2022, Terranova et al 2023, Tonegawa et al 2018, Wang et al 2006, Yamamoto et al 2021).

Automated cell detection algorithms are used to increase efficiency and minimize observer bias (Franceschini et al 2020, Guzowski et al 2005, Moffitt et al 2018, Renier et al 2016, Sundquist & Nisenbaum 2005). A variety of automated cell detection methods have been proposed and used for decades (Bal et al 2020, Benali et al 2003, Carpenter et al 2006, Molnar et al 2016, Sekar et al 2021, Thomas & John 2017, Xing & Yang 2016). However, their performance is frequently demonstrated with high-contrast images with a clear background, such as images of a cultured sample or a brain region with homogenous structure. In actual analysis of fixed brain tissue image, identification of IEG-positive cells is often challenging due to inhomogeneous background arising from uneven staining, localized out-of-focus signals, and autofluorescence of complicated brain-structures (Chen et al 2006, Costantini et al 2021, Fu et al 2024, Wagnieres et al 1998). Global intensity thresholding and gradient/edge-detection are the simplest identification methods (Loukas et al 2003, Wadduwage et al 2018, Wahlby et al 2004), but they are prone to erroneous detections due to complex background (Jones et al 2006, Ofir et al 2020, Waters 2009). Feature extraction methods perform detection and segmentation of cells by finding similarities in features such as intensity and morphology, yet appropriate background correction is still needed as a preprocessing step because it impairs intensity measurements and cellular feature identification (Bray et al 2016, Thomas & John 2017). Machine learning based methods have recently been started to be used for cell-detection (Berg et al 2019, Clissa et al 2024, Pachitariu & Stringer 2022, Schmidt et al 2018, Sommer et al 2011, Stringer et al 2021) and background-subtraction (Buggenthin et al 2013). However, their performance depends on training by researchers or pretrained models, and may suffer from overfitting problem (Laine et al 2021, Moen et al 2019, Nichols et al 2019). Also, uneven background still affects cell detection quality in the machine-learning based methods (Kleinberg et al 2022). Thus, the search for a robust, efficient, and practically usable automated cell detection method is still ongoing.

Many methods have been proposed to suppress background effects for automatically identifying cells, assuming smooth-gradient background such as those arising from non-uniform illumination and aberration (Leong et al 2003, Model & Burkhardt 2001, Yang et al 2015, Zinchuk & Grossenbacher-Zinchuk 2009). Spatial frequency filtering separates high spatial frequency signals from low frequency background (Hupfel et al 2021, Sandison & Webb 1994, Xu et al 1994), but it is less powerful when the background contains high frequency components. Rolling-ball algorithm is a represented method to subtract heterogenous background (Ho et al 2011, Moeyaert et al 2018, Sternberg 1983), yet it has a risk to cause artifact depending on the parameter and sample structure (Gassmann et al 2009, Sticker et al 2020). Adaptive local thresholding determines multiple thresholds for each local regions of an image (Neerad et al 2011, Riccio et al 2019, Sezgin & Sankur 2004), which is similar to Gaussian blur filtering in terms of estimating background based on local pixels (Leong et al 2003). However, these method can fail when signals, or labeled cells, have excessively high intensities or when cells are dense so that they occupy large parts of the local region of the image (Goh et al 2018, Kittler & Illingworth 1985, Lee & Park 1990).

In this study, we proposed an automated cell detection algorithm with a simple background-subtraction method through improving the conventional adaptive thresholding. To prevent erroneous background assumption, excessive intensity pixels were excluded from background assuming calculation by intensity histogram of the image. Cells were then detected by applying intensity thresholding to the background-subtracted image. The performance of the proposed method was assessed with the images of c-Fos and NeuN stained sections in the prefrontal cortex (PFC) and the hippocampal dentate gyrus (DG) of mouse brains. We demonstrated the proposed algorithm’s ability to investigate stimulation-induced increase in IEG-positive cells, identify cells colocalized with different neural markers, and track changes in IEG expression level.

## 2. Materials and methods

### Animals

All procedures relating to mouse care and experimental treatments conformed to NIH and Institutional guidelines, and were conducted with the approval of the UT Southwestern Institutional Animal Care and Use Committee (IACUC). Total 18 male C57BL/6J mice between 8–16 weeks old were used. Mice were group housed with littermates (2–5 mice per cage) until 1 day before experiments in a 12-hour (6am-6pm) light/dark cycle. Mice had ad libitum access to food and water.

## Sample preparation and Imaging

We prepared brain section samples with two different conditions: home cage (HC) and Contextual fear conditioning (CFC) groups (six animals for each group, three to six sections for each brain region were obtained per animal).

### Home cage

Mice were separated into individual cages one day prior to sampling. The mice were deeply anesthetized with a ketamine (75 mg/kg)/dexmedetomidine (1 mg/kg) cocktail and transcardially perfused with 4% paraformaldehyde in PBS (PFA). Brains were removed and post-fixed in 4% PFA at 4°C for 24 hours.

### Contextual fear conditioning

Mice were separated into individual cages 1 day prior to the conditioning. Foot shock was provided based on a previous reported contextual fear conditioning paradigm (Terranova et al 2022). The shock apparatus consists of a chamber (24 cm W × 20 cm D × 20 cm H) (Med Associates). Each mouse was placed in the stimulation chamber for a 3-minute habituation period. Subsequently, during a 3-minute shock period, the mouse received 0.75 mA, 2-second foot shock with a 58-second shock interval between shocks, for a total of three shock trials. After the stimulation, the mouse was returned to the home cage. One hour later, the mouse was deeply anesthetized with a ketamine/dexmedetomidine cocktail and transcardially perfused with 4% PFA. Brains were removed and post-fixed in 4% PFA at 4°C for 24 hours. The chamber was cleaned before starting this protocol with each mouse.

### Immunohistochemistry and Imaging

Post-fixed brains were sectioned at 60 μm thickness using a vibratome (Leica VT100S). For immunohistochemistry (IHC), tissue sections were incubated in 0.03% Triton-X PBS (PBS-T) with 5% normal donkey serum (NDS) for 30 minutes. Subsequently, the sections were incubated with primary antibodies diluted in PBS-T with 5% NDS for 2 overnight at 4°C: chicken anti-NeuN (1/1000, Millipore Sigma, ABN91) and goat anti-cFos (1/1,000, Santacruz, sc-52-G). After rinsing with PBS (3 × 5 min), tissue sections were incubated with secondary antibodies in PBS-T with 5% NDS for 2 hours at room temperature: donkey anti-chicken AlexaFluor405 (1/500, Jackson ImmunoResearch Labs) and donkey anti-goat AlexaFluor633 (1/500, Jackson ImmunoResearch Labs). Tissue sections were then washed in PBS (3 × 5 min) and mounted on glass slides with VECTASHIELD antifade mounting medium (Vector Laboratories). All fluorescence images (0.624 µm/pixel) were acquired with Zeiss LSM800 confocal microscope using 10× objectives (NA: 0.45), under consistent imaging condition for laser intensity, pinhole size, scan speed, and image acquisition gain. Images were exported to 8-bit images using the Zen Blue software (Zeiss).

#### Analysis

c-Fos- and NeuN-positive cells were detected in the prelimbic (PL) part of medial prefrontal cortex (mPFC), and the granule cell layers of dorsal hippocampal dentate gyrus (DG). The boundaries of the brain regions were determined according to the Allen Brain Reference Atlas (Allen Institute for Brain Science 2004). From bregma, PL was determined in the sections at +1.845 to +1.42 mm, while dorsal DG was at -1.655 to -2.255 mm. Regions of interest (ROI) for each brain section were created manually using ImageJ (Schindelin et al 2012). The images of each channel (NeuN and c-Fos) and the ROI information were imported into MATLAB R2023a (MathWorks) for further analysis.

### Automated cell detection

The custom codes were written in MATLAB with Image Processing Toolbox (Fig. S1). The automated cell detection process consists of five steps: (1) pre-processing, (2) background assumption and subtraction, (3) thresholding, (4) denoising and smoothing, and (5) cell-segmentation (Fig. 1). The key step of the proposed method is (2) background assumption and subtraction, where the background is assumed based on local average intensities of the image avoiding strong intensity pixels which were determined from the histogram of the image. Intensity thresholding was applied after background subtraction. Our approach is similar to standard adaptive thresholding, but has a difference in avoiding putative signal pixels in the calculation. The processes of each step are as follows (Fig. 1C).

**Fig. 1.**
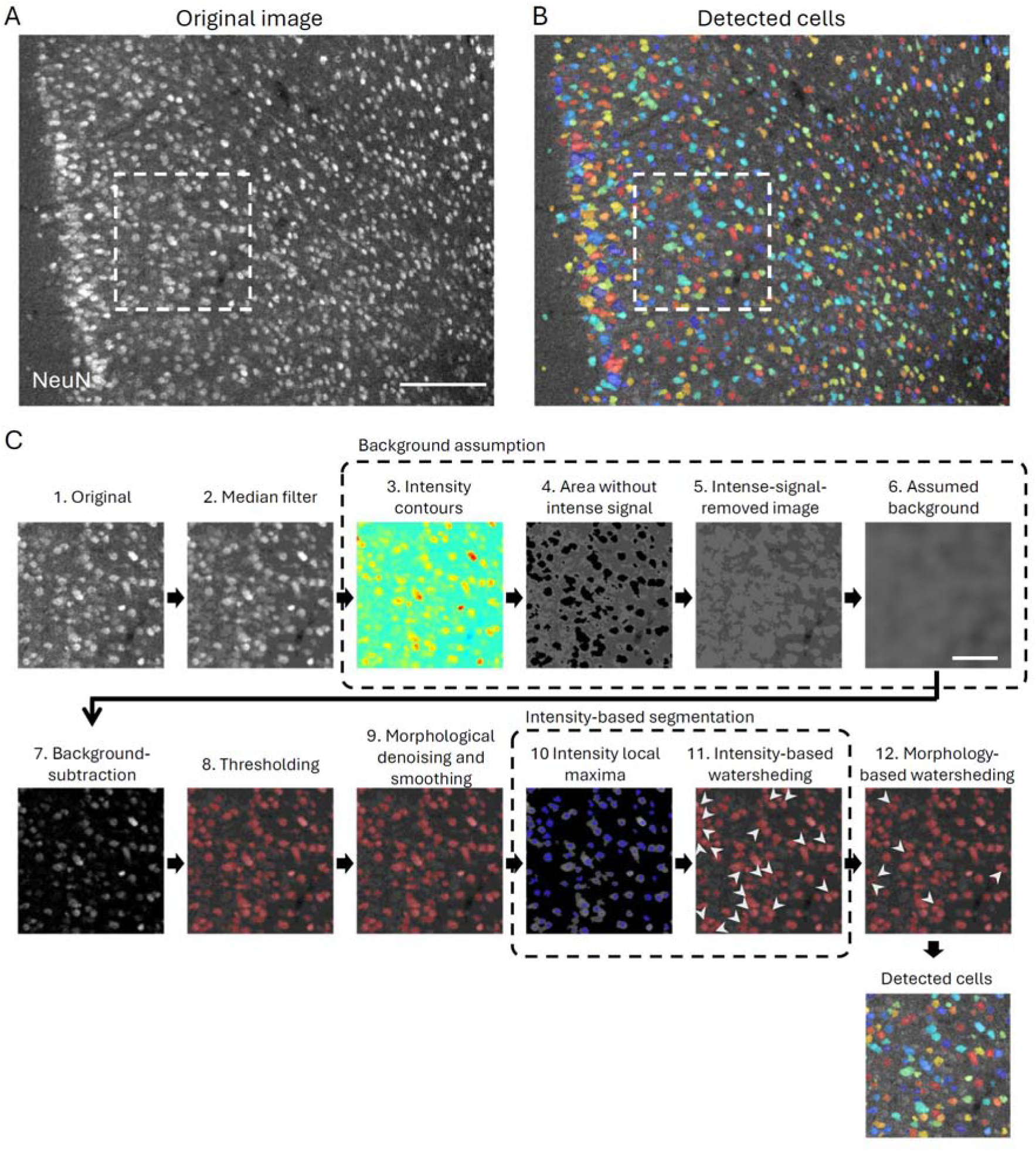
Protocol of background assumption and cell detection. (A) Original image of NeuN stained cells in PFC. (B) Result of automated cell detection (colored). White dashed line in (A) and (B) indicate the area shown in (C). (C) Overview of the background assumption and cell detection protocol. The fluorescent labeled cell image (1) was median filtered as a preprocessing step (2). To assume the background, twenty intensity contours were calculated by Otsu’s method (3), with a contour occupying >80% was used as pixels for background assumption (4). Pixels inside of the drawn contour were filled with the same intensity as the contour (5), followed by blurring using a moving average filtering of 31-pixels to create the assumed background pattern (6). The assumed background was then subtracted (7). Then, signals were thresholded based on intensity (8) (see Methods). The thresholded image was denoised/smoothed by morphological processing (9), and then cells were segmented by watersheding algorithm (10-12; white arrows indicate segmented cells). Scale bars, 200 µm (A) and 100 µm (C).

(1) As a pre-processing, the imported images were converted to the 8-bit gray-scale images, and applied median filter (NeuN image: 7×7 pixels; c-Fos image: 11×11 pixels). (2) Background pattern was assumed and subtracted. First, we determined the intense signal pixels in an image by drawing total twenty intensity contours using Otsu’s method (Otsu 1979), which is implemented in MATLAB Image Processing Toolbox (*multithresh.m*). A contour which covers more than 80% (for NeuN image) or 95% (for c-Fos image) of the image was selected for further calculation. The areas insides of the contours were filled with the intensity of neighboring pixels, and the background pattern was assumed by smoothing with moving average filtering (31×31 pixels). The assumed background was then subtracted from the median-filtered image. (3) Thresholding was conducted on the background-subtracted image with two criteria. The first criterion is standard-deviation based global thresholding, defined as T = k · CJ, where T is the threshold, σ is standard deviation intensity of the background-subtracted image, and k is coefficient parameter (k = 2 for NeuN+ and k = 5 for c-Fos+ cell detections). The second criterion is histogram-based global thresholding, using Otsu’s method in the background-subtracted image. Pixels with intensities below zero value in the background-subtracted image were substituted with zeros, then twenty intensity contours were drawn by Otsu’s method. A contour covering more than 80% of the image area was selected as an additional threshold. The motivation of the second criterion is to reduce false-positive detection that may arise solely using the first criterion, because the standard-deviation based thresholding can cause misclassification when the most part of the image is occupied with background (Yan et al 2005). Pixels that met both thresholds were classified as signals. (4) Thresholded signals were denoised and smoothed with morphological operations (Serra & Vincent 1992) using a binarized image of the thresholded data. First, the image was denoised by erosion with a diamond-shaped element with five pixels distance followed by reconstruction operation. Then, the image was smoothed by morphological closing by a diamond-shaped element with two pixels distance. In addition, signals whose areas are smaller than 50 pixels are ignored and removed for further processing. (5) To segment closely positioned cells, we employed a watershed algorithm (Malpica et al 1997, Meyer 1994) with two steps. The first step is based on the regional intensity maxima of the image. Watershed delineation was performed after finding regional maxima, and the signals were segmented along the created lines. In the second step, the signals were further segmented based on cell morphology. Tiny local minima were removed using *imextendedmin.m* function of MATLAB to avoid over-segmentation (Ismail et al 2016), and then watershed delineation was performed on the image to segment cells. The segmented signals are counted as detected cells if their areas exceeded 50 pixels. Maximum intensities within the detected cells were used to investigate IEG expression level. Cells that share more than five pixels between different neural markers are considered double-labeled. The custom MATLAB code used in this study is available at https://github.com/HisayukiOsanai/CellDetection.

To evaluate the detection quality of the automated method, manual cell detection was performed using ImageJ by an experimenter who was blinded to the result of the automated cell detection. The false positive detection rate, or Precision, was calculated as Precision = TP / (TP + FP), where true positive (TP) and false positive (FP) were checked manually after the automated detection with randomly selected section images. In addition, the Auto-Manual match rate, or Sensitivity, was calculated to evaluate the ratio that cells manually detected were also detected in the automated algorithm. To Compare with the manual detection performed by an experimenter who was blinded to the automated detection results, the Auto-Manual match rate was calculated as Match / (Match + Eye_Only), where Match indicates the number of cells identified both by the automated detection and manual detection, and Eye_Only indicates the number of cells identified manually but not in the automated detection. Furthermore, Sensitivity-increase rate was calculated to evaluate the ratio that cells were not detected manually by the experimenter but detected in the automated algorithm, as Auto_Only / (Match + Auto_Only) where Auto_Only indicates the number of cells that were not identified manually but detected in the automated algorithm. For comparison, Rolling ball algorithm implemented in ImageJ (Schindelin et al 2012) was performed with a parameter radius of 22.3 µm (Fig. S3). Comparison with manual detection was performed with sections in which cell signal was detected.

### Statistics

Statistical analyses were performed using MATLAB. We used a one-tailed student’s t test (Fig. 5F, 5G, 6F, 6G) or a two-tailed Wilcoxon signed rank test (Fig. 6H). p < 0.05 was assumed to be statistically significant. ∗ indicates p < 0.05, ∗∗ indicates p < 0.01, and ∗∗∗ indicates p < 0.001. Bar plots and error bars represent means and standard errors.

## 3. Results

### 3.1. Assumption of background pattern and performance of automated cell detection

The proposed automated cell detection method was first tested on confocal images with relatively homogeneous background in PFC (Fig. 1-3) in which cells were stained with anti-NeuN and anti-c-Fos antibodies. Background assumption was performed with blur filtering avoiding image pixels of excessive intensities. The assumed background was then subtracted for further thresholding, denoising and cell segmentation processes (Fig. 1C). Next, we assessed the accuracy of our proposed automated cell-detection algorithm by evaluating false positive rate and by comparing with manual detection. The false positive detection, assessed manually after the automated detection, was very low (Precision > 0.98) in NeuN stained images in the PFC (Fig. 2A-D). The false positive detections were mostly due to cell over-segmentation in watersheding processes. In addition, the cell detection performance was compared to manual detection by an experimenter who was blinded to the automated detection result (Fig. 2E-G). The automated detection identified more than 90% of the cells that were identified manually (Fig. 2E, Auto-Manual match rate > 0.90) with increased sensitivity (Fig. 2F, Sensitivity-increase rate = 0.29 ± 0.07), indicating our proposed method performs with high sensitivity. The performance was further tested in c-Fos stained images (Fig. 3), similarly comparing the results of automated and manual detection (Fig. 3A-C). The precision rate was comparably high (>0.99; Fig. 3D). While NeuN-positive cell intensities were relatively similar between cells (Fig. 1A, 2A), the intensities of c-Fos-positive cells were largely variable between cells that led to operator bias of manual detection; cells of similar intensities tended to be counted when isolated from other cells, but were often overlooked when surrounded by higher intensity cells (Fig. S2B, C), resulting in difference between the automated and the manual detection performance (Auto-Manual match rate = 0.70 ± 0.04; Sensitivity-increase rate = 0.32 ± 0.06) (Fig. 3E-G). The results demonstrated that our proposed detection algorithm has high precision and helps reduce operator bias in the manual detection.

**Fig. 2.**
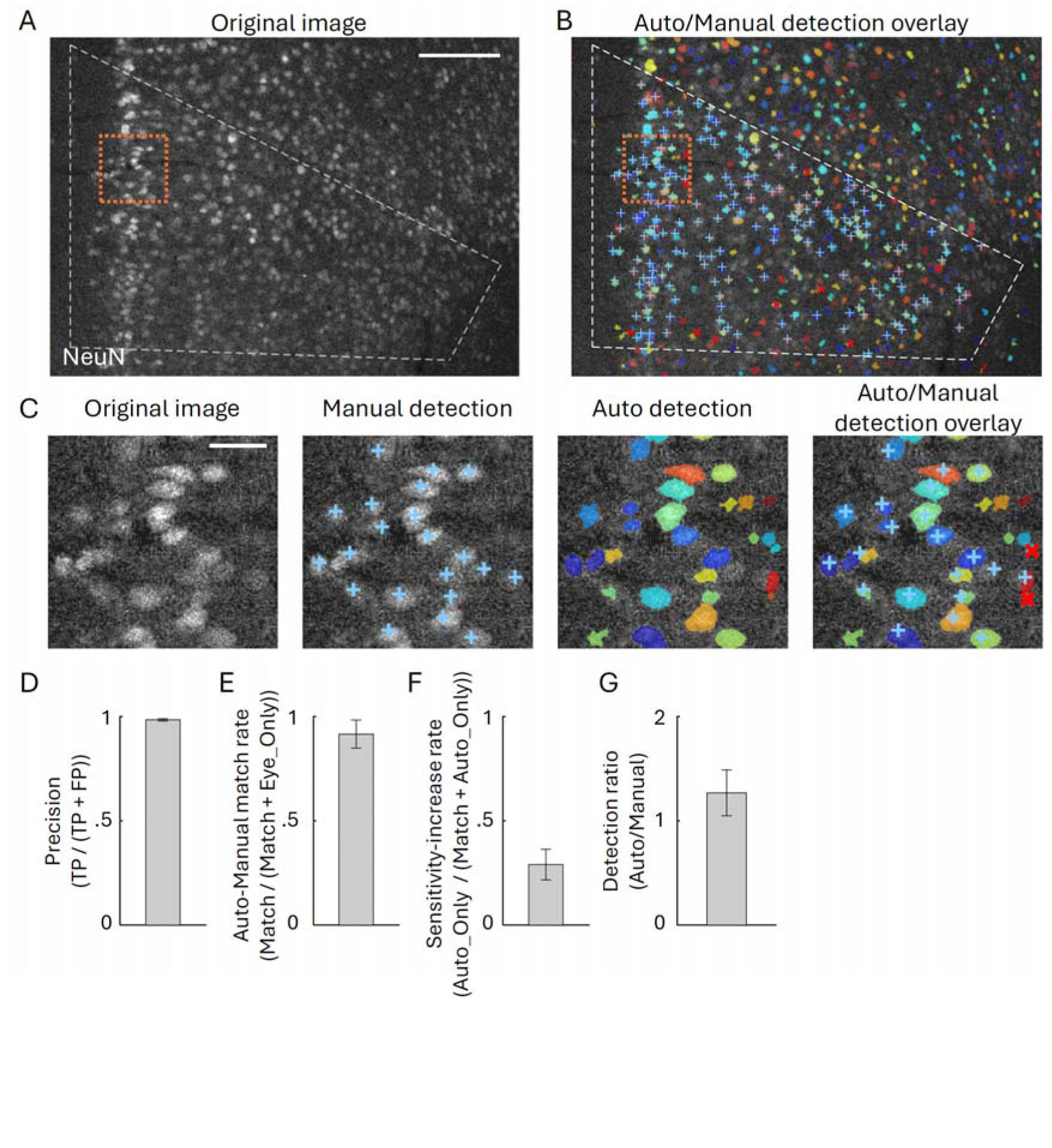
Measurement of cell detection accuracy comparing between automated and manual detection in NeuN staining sections. (A) Original image of NeuN stained cells in the PFC. White dashed line indicates the ROI of the PL area. Orange dotted line indicates the area shown in (C). (B) Results of automated cell detection (colored), along with manual detection within the ROI by an experimenter who was blinded to the automated cell detection result (light-blue ‘+’ mark). The red ‘×’ marks indicate the manually verified false-positives in the automated detection, that were mostly due to cell over-segmentation by whatersheding. (C) Magnified views of the original image, manual detection, automated detection, and their overlay. (D-G) Quantification of the detection precision and matching/mismatching ratio compared between automated and manual detection (see Methods). (D) Precision of the automated detection, which indicates false positive ratio in the automated cell detection per total detected cells (n = 3 sections). Precision = TP / (TP + FP), where true positive (TP) and false positive (FP) were checked manually after the automated detection. (E) Match rate between automated and manual detection, indicating the ratio that cells manually detected were also detected in the automated algorithm (n = 3). Auto-Manual match rate = Match / (Match + Eye_Only), where Match indicates the number of cells identified both by the automated and manual detection, and Eye_Only indicates the number of cells identified only by manually. The manual detection was conducted by an experimenter who was blinded to the automated detection results. (F) Sensitivity increase rate, indicating ratio that cells were not detected manually by the experimenter but detected in the automated algorithm. Sensitivity-increase rate = Auto_Only / (Match + Auto_Only), where Auto_Only indicates the number of cells that were not identified manually but detected in the automated algorithm. (G) Ratio of the number of automated and manually detected cells (n = 3). Scale bars, 200 µm (A) and 40 µm (C).

**Fig. 3.**
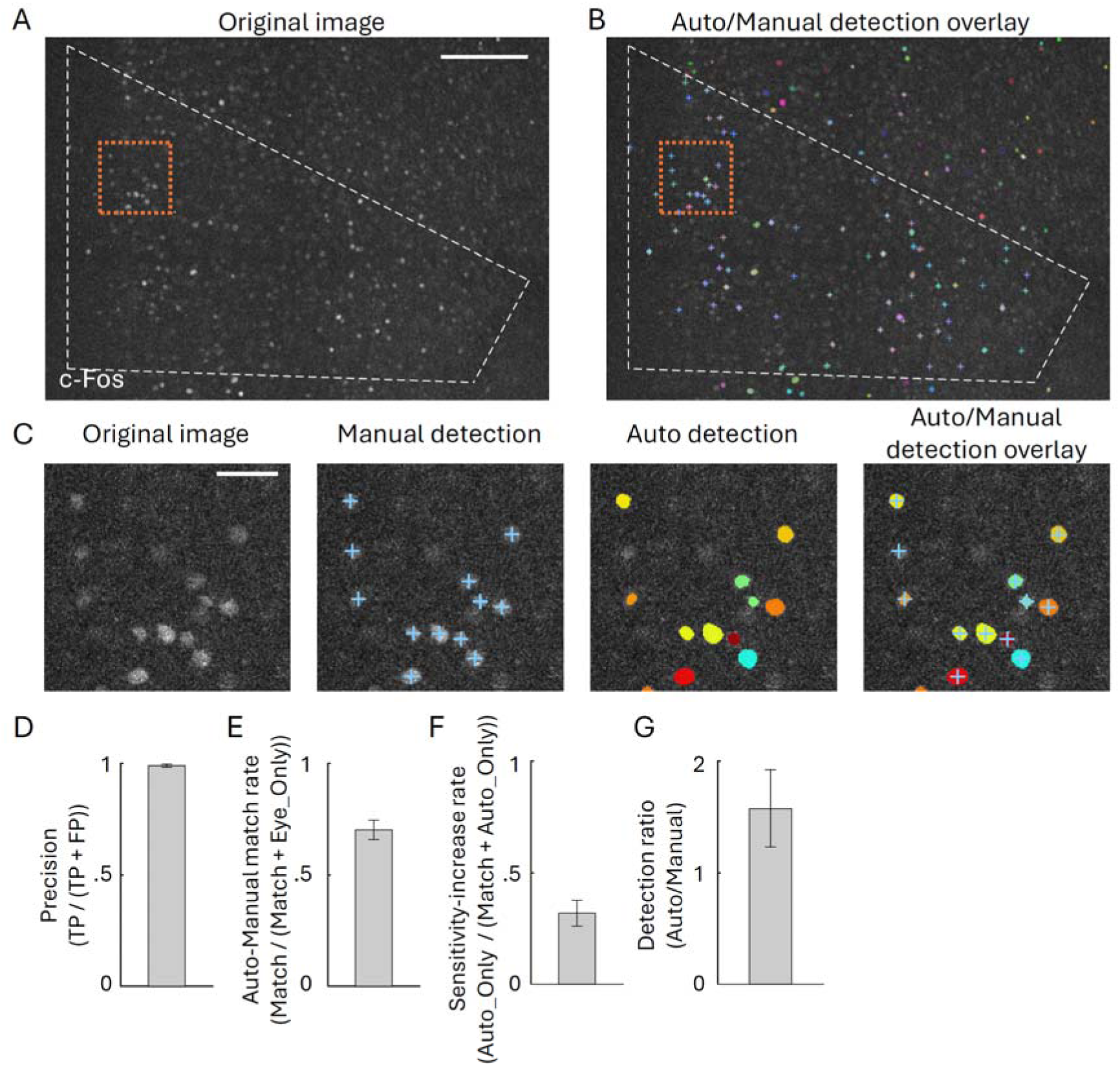
Comparison between automated and manual cell detection in c-Fos stained sections. (A) Original image of c-Fos stained cells in the PFC, same brain section shown in Fig. 2. White dashed line indicates the ROI of the PL area. Orange dotted line indicates the area shown in (C). (B) The results of automated cell detection (colored), and manual detection within the ROI by an experimenter who was blinded to the automated cell detection result (light-blue ‘+’). (C) Magnified views of the original image, manual detection, automated detection, and their overlay. (D-G) Quantification of the detection precision and matching/mismatching ratio compared with manual detection, similarly to Fig. 2D-G. Precision of the automated detection (E; n = 6 sections), match rate between automated and manual detection (F; n = 20), sensitivity increase rate (F; n = 20), and ratio of the number of automated/manually detected cells (G; n = 20). Scale bars, 200 µm (A) and 40 µm (C). Images shown in (A-C) are representative images from Contextual fear conditioning (CFC) group, and data shown in (D-G) are both from home cage (HC) and CFC groups.

Next, we tested our proposed cell-detection algorithm on images with inhomogeneous background in DG (Fig. 4). Our algorithm was able to assume the background pattern by avoiding excessive intensity pixels, resulting in the image in which the inhomogeneous background was effectively removed (Fig. 4A-E, Fig. S3B). The automated detection of c-Fos-positive cells showed a low false positive rate (Precision > 0.97; Fig. 4F-H). Compared to manual detection results, the automated detection in DG identified more than 70% of the manually identified cells (Fig. 4F, G, I, Auto-Manual match rate = 0.73 ± 0.03; Fig. 4J, Sensitivity-increase rate = 0.08 ± 0.02), similarly to the results in the PFC (Fig. 3E). Additionally, compared to the conventional Rolling ball algorism, our method has the advantage of minimizing the risk of artifacts in densely packed cells (Fig. S3). These results indicate that our method can obtain a background-free image for further automated cell detection process from an image with inhomogeneous background pattern.

**Fig. 4.**
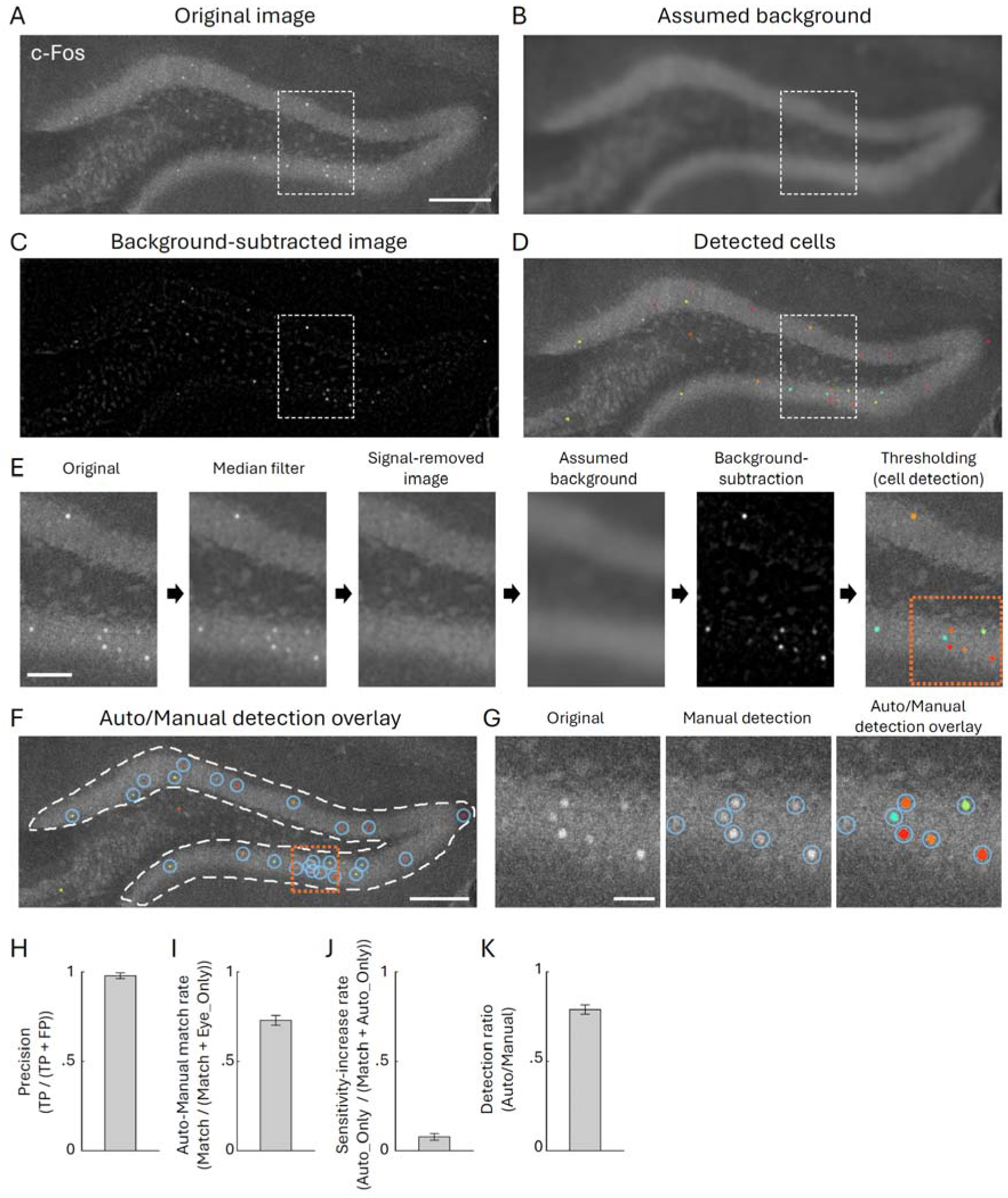
Cell detection on inhomogeneous background image in DG. (A) Original image of c-Fos stained cells in the DG, with observable background patterns along the granule cell layer. White dashed line indicates the area shown in (E). (B) Assumed background pattern. (C) Background-subtracted image. (D) The automatically detected cells (colored). (E) Simplified overview of the automated cell detection process under inhomogeneous background, demonstrating background assumption and subtraction. The detailed process is described in Fig. 1C and Methods. Orange dotted line indicates the area shown in (G). (F-H) Comparison with manual detection, similarly to Fig. 2 and 3. (F) Results of automated cell detection (colored) and manual detection within the ROI (light-blue circles). Orange dotted line indicates the area shown in (G). (G) Magnified views of the orange line area in E and F, original image, manual detection, and their overlay of the automated/manual detection. (H) Precision of the automated detection (n = 6 sections). (I) Match rate between automated and manual detection (n = 30). (J) Sensitivity-increase rate (n = 30). (K) Ratio of the number of automated and manual detected cells (n = 30). Scale bars, 200 µm (A, F), 100 µm (E), and 40 µm (G). Images shown in (A-G) are representative images from HC group, and data shown in (H-K) are both from HC and CFC groups.

### 3.2. Detection of increased c-Fos-positive cells after stimulation

To validate the utility of our method, we performed the detection of c-Fos-positive cells in the PFC and DG of homecage (HC; Fig. 5A, C) and contextual fear conditioning groups (CFC; Fig. 5B, D). The detected cell number was correlated between the results of automated and manual counting (Fig. 5E). Increased number of c-Fos-positive cells was observed in the automated detection, similar to the manual detection in the PFC (Fig. 5F; Manual: Cell density = 3.01 ± 1.60 vs. 140.4 ± 21.4 /mm^2^, p = 0.0012; Auto: 16.3 ± 11.2 vs. 121.2 ± 15.0 /mm^2^, p < 0.001) and DG (Fig. 5G; Manual: Cell density = 108.1 ± 7.4 vs. 163.2 ± 13.5 /mm^2^, p = 0.0013, Auto: 82.9 ± 9.6 vs. 129.1 ± 9.4 /mm^2^p = 0.0011), consistent with many other studies (Beck & Fibiger 1995, Morrow et al 1999). This confirms our method is capable of detecting stimulation-induced changes in the number of IEG-expressing cells.

**Fig. 5.**
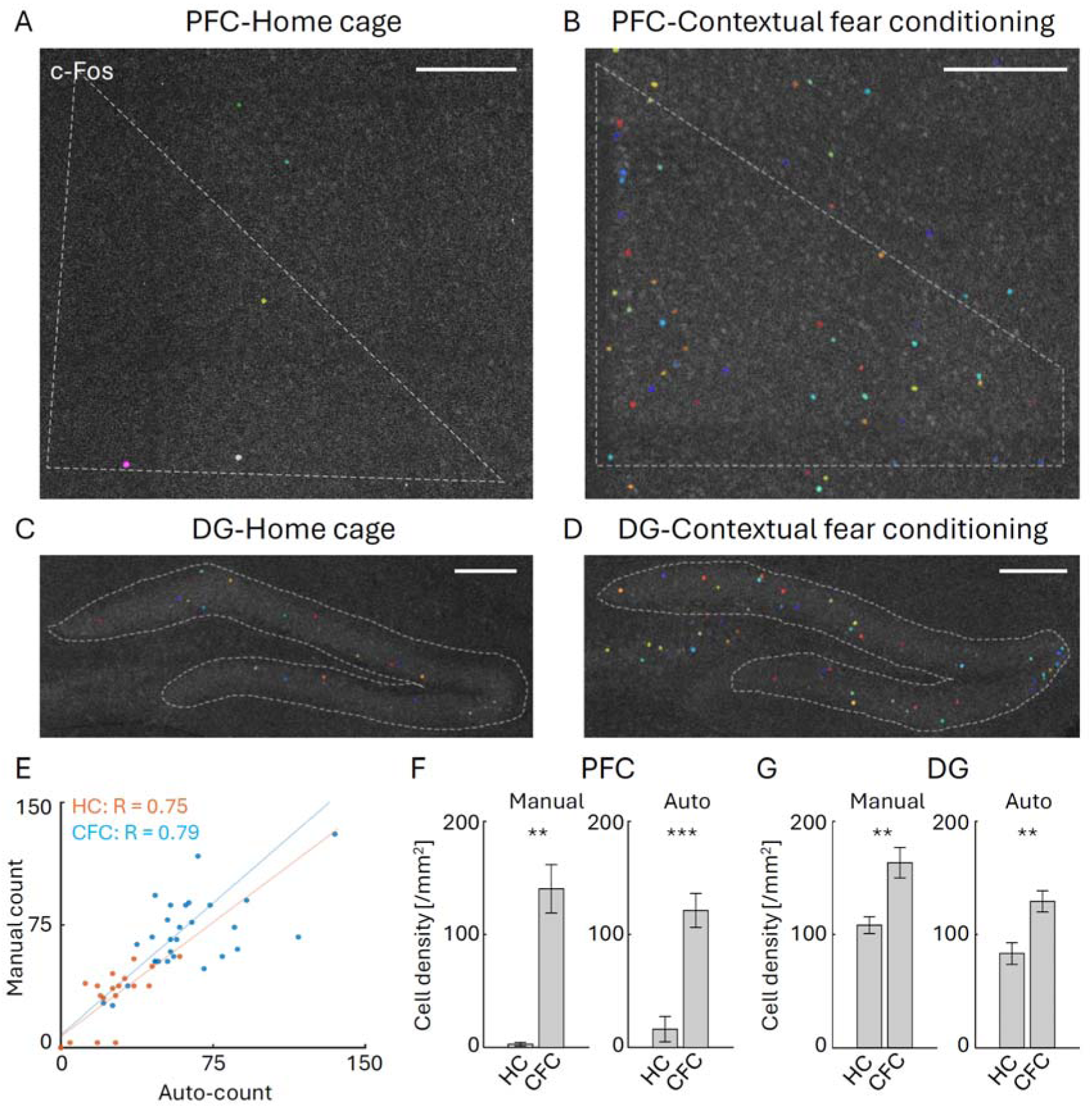
Detection of increased c-Fos positive cells by CFC stimulation. (A, B) Automated detection of c-Fos stained cells in the PFC for home-cage (HC) (A) and Contextual fear conditioning (CFC) groups (B). White dashed lines indicate ROI of PL. (C, D) Automated detection of c-Fos stained cells in the DG for HC (C) and CFC groups (D). White dashed lines indicate ROI of the DG granule cell layer. (E) Correlation of automatically and manually detected cell number, with R indicates Peason correlation coefficient (p<0.001 for both HC and CFC groups) (HC: n = 18, CFC: n = 34 sections). (F) Cell densities of manually (left) and automatically detected cells (right) in the PFC for HC (n = 5) and CFC groups (n = 17). (G) Cell densities of manually (left) and automatically detected cells (right) in the DG for control (HC, n = 13) and CFC groups (n = 17). Scale bars, 200 µm.

**Fig. 6.**
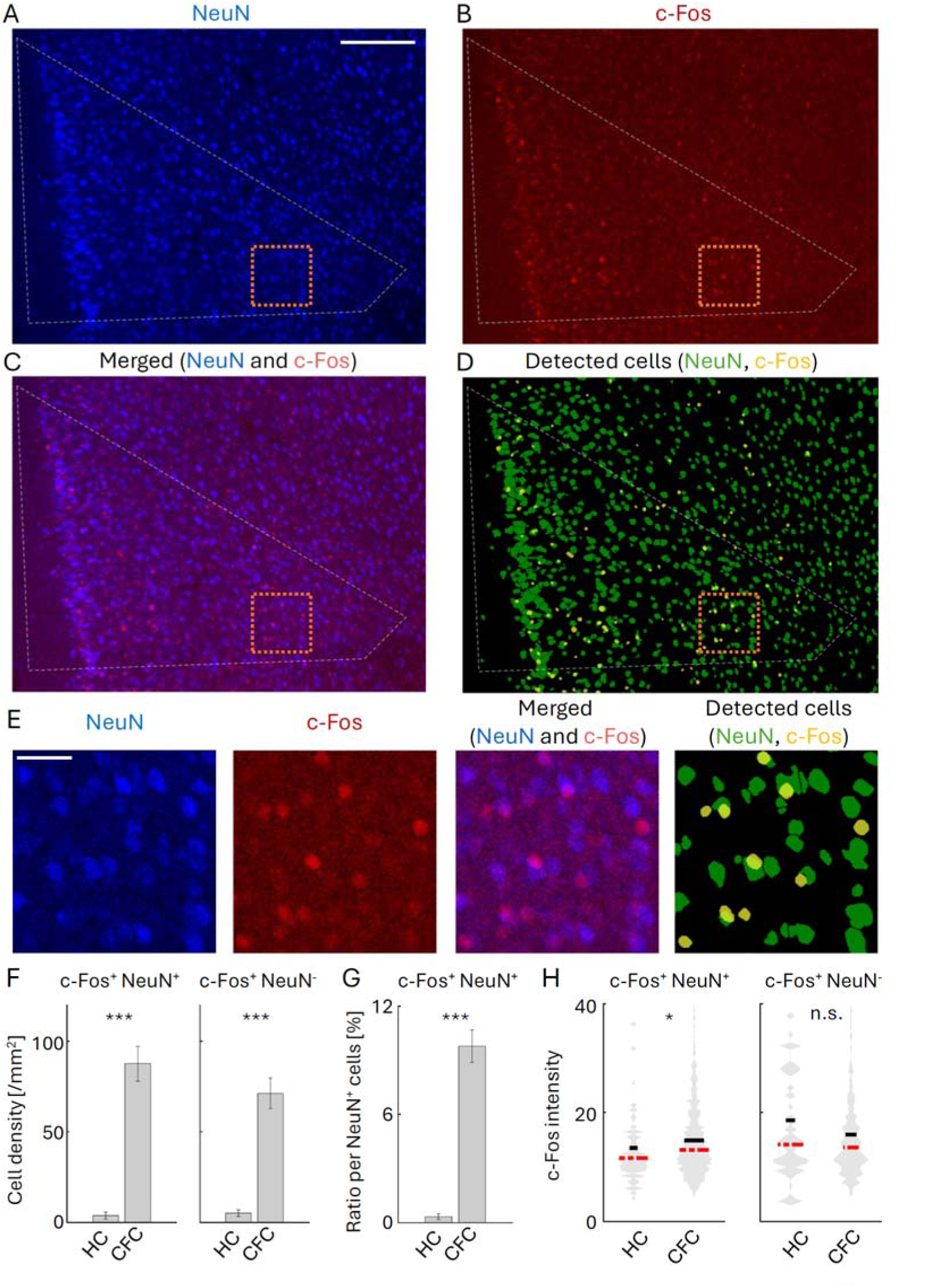
Detection and quantification of c-Fos positive cells with neural marker. (A-C) Images of NeuN and c-Fos co-stained cells in PFC for CFC group. (A) NeuN, (B) c-Fos, and (C) merged images. White dashed lines indicate ROI of the PL. (D) Automated cell detection image. Green: NeuN; Yellow: c-Fos. Orange dotted line indicates the area shown in (E). (E) Magnified views of NeuN, c-Fos, merged, and detected cell images. (F) Cell density within ROI of c-Fos+ NeuN+ (left) and c-Fos+ NeuN- cells (right), in HC (n = 23 sections) and CFC (n = 25 sections) groups. (G) Percentage of c-Fos+ NeuN+ cells per all NeuN+ cells. (H) Intensity of c-Fos signals of c-Fos+ NeuN+ cells (left, HC: n = 73 cells, CFC: n = 780 cells) and of c-Fos+ NeuN- cells (right, HC: n = 83 cells, CFC: n = 715 cells). Red dashed and black solid line indicate median and mean values, respectively. Scale bars, 200 µm (A) and 40 µm (E).

### 3.3 Detection of double-labelled cells and quantification of IEG expression level

Furthermore, we demonstrated the application of our method for double-labeling examination. The co-localization of c-Fos and NeuN signals was imaged in the PFC (Fig. 6A-C, E) and detected automatically (Fig. 6D, E). Within c-Fos-positive cells, we separated the cells into NeuN-positive and NeuN-negative groups. Our method revealed an increase in cell number following CFC both in the c-Fos+ NeuN+ group (HC vs. CFC, Cell density = 3.85 ± 1.91 vs. 87.64 ± 9.50 /mm^2^, p < 0.001; Cell number in percentage per all NeuN-positive cells: 0.33 ± 0.16 vs. 9.76 ± 0.91 %, p < 0.001), and in the c-Fos+ NeuN- group (HC vs. CFC, Cell density = 5.21 ± 1.86 vs. 71.33 ± 8.35 /mm^2^, p<0.001) (Fig. 6F, G). On the other hand, while we observed that c-Fos intensity was increased by CFC in the c-Fos+ NeuN+ group (p = 0.019), the intensity was not significantly changed in the c-Fos+ NeuN- group (p = 0.65) (Fig. 6H). This suggests that c-Fos expressing levels are differently regulated between NeuN-positive and NeuN-negative cells. Overall, our proposed method is useful for efficient and unbiased analysis of fluorescent labeled cells in images with inhomogeneous backgrounds.

## 4. Discussion

Although many methods have been proposed for cell detection in fluorescent labeled images, not all of them are assuming working on highly inhomogeneous background. While there are efforts to experimentally suppress background intensity (Sun et al 2017, Weiss et al 2021), it is difficult to remove the background completely, thus analytical background removal is required in many cases. Several background assumption methods utilize three-dimensional (Biggs 2010, Yang et al 2015) or time-lapse information (Peng et al 2017), but such information are not always available. In this study, we propose an algorithm to assume and subtract inhomogeneous background in two-dimensional fluorescent image by avoiding excessive intensity pixels for blur calculation. This approach overcomes limitations in the conventional adaptive local thresholding techniques. The background-subtracted images were then effectively used for automated cell detection.

Our automated cell detection achieved a high precision rate. Since the errors in our method were mostly due to over-segmentation of cells, combining our method with other state-of-art cell-segmentation protocols (Molnar et al 2016, Schmidt et al 2018, Wang 2019, Xing & Yang 2016, Zhang et al 2021) will further increase detection accuracy on images with inhomogeneous background structure. Because our algorithm does not have any iterative calculation except watersheding segmentation process, our method has an advantage in fast computation so that it is easy to combine with other cell segmentation or identification methods. Thus, our background-subtraction method can serve as a pre-processing protocol for various other cell segmentation techniques.

In the demonstration of application for double-labeling experiments, we found that the number of the c-Fos+/NeuN+ cells increased by CFC similarly to the c-Fos+/NeuN- cells. On the other hand, the increase in c-Fos intensity was not observed in the c-Fos+/NeuN- cells, in contrast to the c-Fos+/NeuN+ cells. Since NeuN express specifically in neuronal cells (Mullen et al 1992), the c-Fos+/NeuN- cells are supposed to be glial cells (Aguilar-Delgadillo et al 2024, Cruz-Mendoza et al 2022, Williamson et al 2024). The increase in c-Fos+ glial cells align with previous findings (Aguilar-Delgadillo et al 2024, Cruz-Mendoza et al 2022). On the other hand, to the best knowledge, the expression level of c-Fos within c-Fos+ glial cells has not been investigated yet. Our results suggest that c-Fos expression level in individual glial cell is strictly regulated compared to those in neuronal cells, once c-Fos is expressed. However, further co-labelling investigation is needed with glial cell markers.

## 5. Conclusions

In this study, we developed an automated cell detection algorithm that effectively subtracts inhomogeneous backgrounds from fixed brain tissue images. The results indicate that the proposed algorithm performs adequately to investigate changes in IEG-expressing cells and their expression levels induced by stimulation. Our method offers significant advantages in providing high-throughput and unbiased analyses of images from regions with complex biological structures that generate inhomogeneous backgrounds. The method will accelerate examination of cell-type specific distribution of memory engram cells in multiple brain regions.

## Ethics Statements

All procedures relating to mouse care and experimental treatments conformed to NIH and Institutional guidelines, and were carried out with the approval of the UT Southwestern Institutional Animal Care and Use Committee (IACUC).

## CRediT authorship contribution statement

**Hisayuki Osanai**: Conceptualization, Methodology, Software, Validation, Formal Analysis, Investigation, Data curation, Writing - Original draft, Writing - Review & Editing, Visualization. **Miari Arai**: Investigation, Data curation. **Takashi Kitamura**: Writing - Review & Editing, Project administration, Supervision, Conceptualization. **Sache K Ogawa**: Writing - Review & Editing, Project administration, Supervision, Conceptualization.

## Acknowledgement

We thank all lab members for their support. This work was supported by grants from the Endowed Scholar Program to T.K., Brain Research Foundation to T.K. (BRFSG-2018-04), Faculty Science and Technology Acquisition and Retention Program to T.K., the Whitehall Foundation to T.K. (2019-05-38), the National Institute of Mental Health to T.K. (R01MH120134, R01MH125916, R01NS138075) and Japan Society for the Promotion of Science to H.O. (201860198 and 202101654). Japan-U.S. Brain Research Cooperative Program: FY2023 to M.A.

## Data availability

Data will be made available on request. MATLAB code is available at https://github.com/HisayukiOsanai/CellDetection.

---

**Fig. S1.**
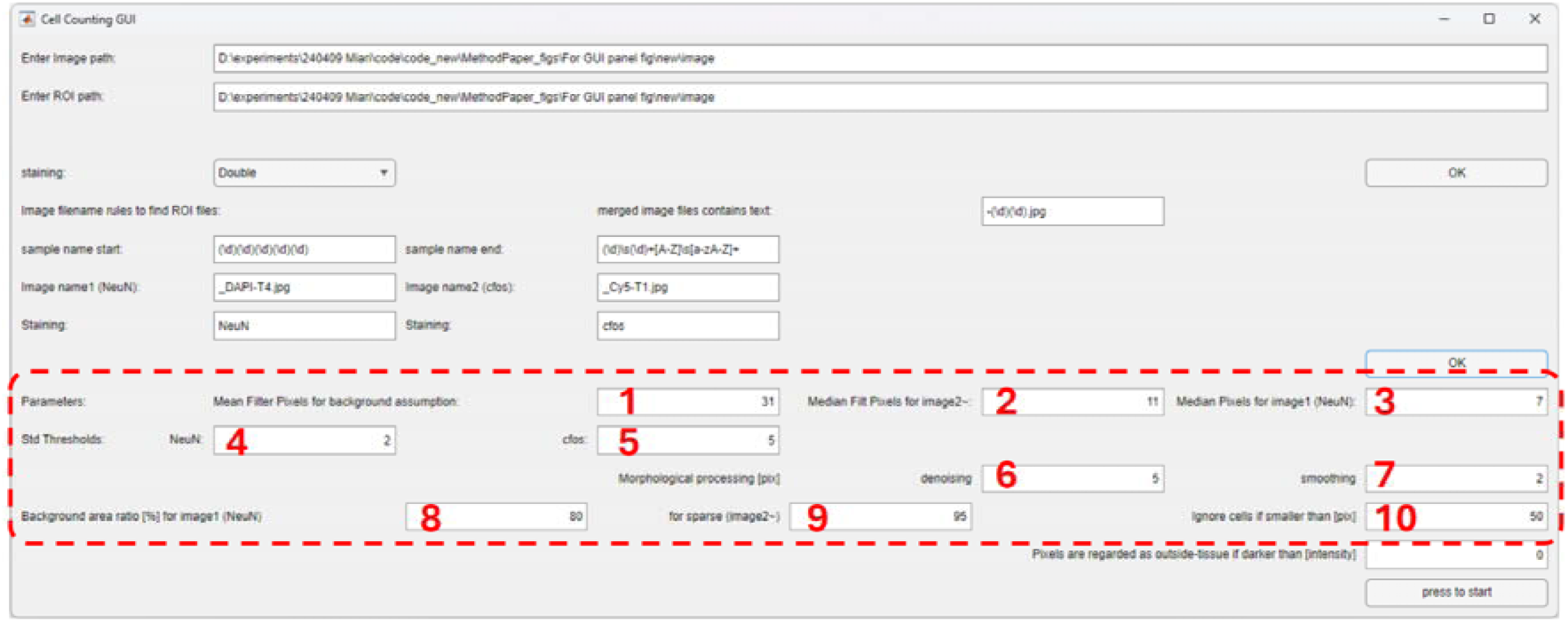
Graphical user interface of automated cell detection. Graphical user interface panel (GUI) of automated cell detection. User-defining parameters for cell-detection include: 1. Pixels used for mean average (blurring) filtering (31 pixels). 2-3. Pixels used for median filtering for c-Fos (2) and NeuN (3) staining image (11 and 7 pixels). 4-5. Standard deviation intensity threshold parameters for NeuN (> 2 * std) and for c-Fos images (> 5 * std). 6-7. Pixels for morphological processing denoising (5 pixels) and smoothing (2 pixels). 8-9. Assumed background area per NeuN staining image (80%) and per c-Fos staining image (95%). 10. Signal areas less than 50 pixels are regarded as noise and not counted as cells.

**Fig. S2.**
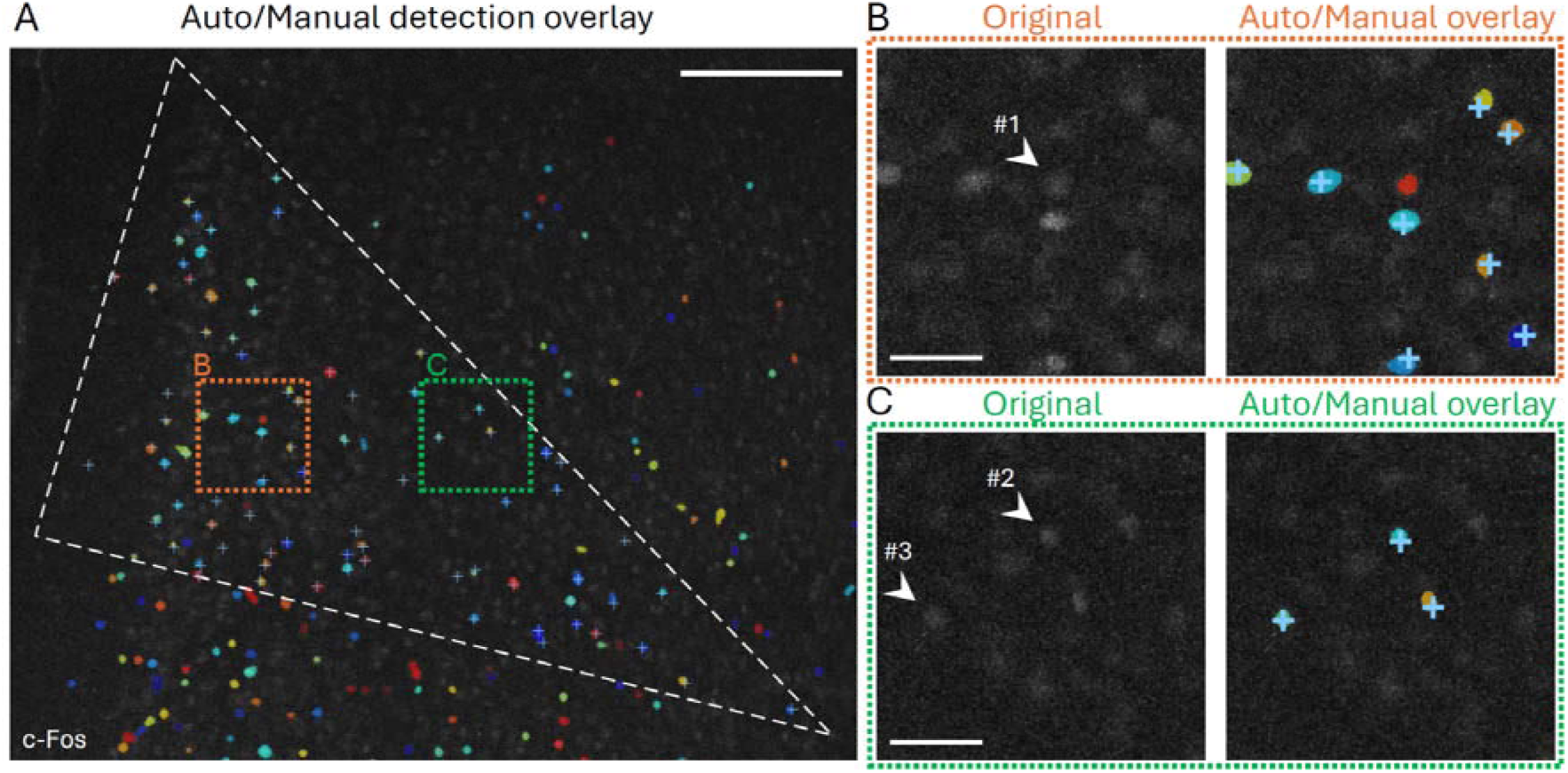
Example of biased cell identification in manual detection. (A) Overlay of automated cell detection (colored) and manual detection within the ROI (light-blue ‘+’) in the c-Fos stained PFC image. (B, C) Magnified views of the original image and automated/manual detection of cell-dense (B) and sparse area (C), which are indicated by orange and green dotted lines in (A). Arrowheads indicate cells with similar fluorescent intensity (cell #1, F = 35; #2, F = 63; #3, F = 31). Cell #2, 3 were identified in the manual detection at the cell-sparse area but cell #1 was overlooked at the cell-dense area, while all of them were identified in an unbiased manner in the automated detection identified. Scale bars, 200 µm (A) and 40 µm (B, C).

**Fig. S3.**
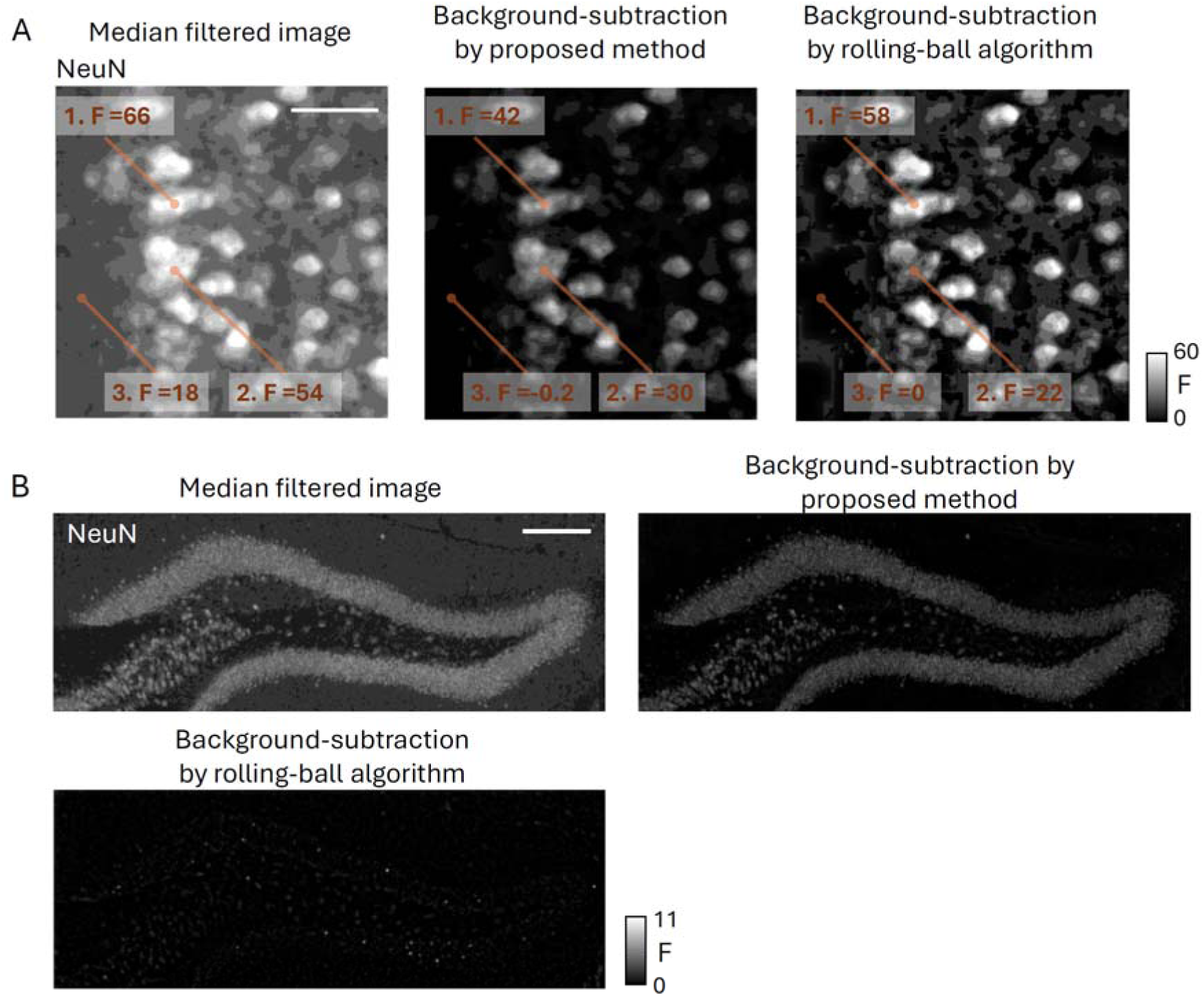
Artifacts caused by Rolling ball algorithm. (A) Example of artifact induced by Rolling ball algorithm. F indicates fluorescent intensity. Left, Median filtered image of NeuN stained cells in PFC. Middle, Background-subtracted image by our proposed method. Right, Background-subtracted image by Rolling ball algorithm. Note that the intensity difference at areas #1 and #2 is 12 in the median filtered image and the background-subtracted image with our method. However, their difference in the Rolling ball algorithm becomes 36, indicating that Rolling ball algorithm subtracted signals too much when cells are closely positioned. (B) Another example of artifact induced by Rolling ball algorithm. Top left, Median filtered image of NeuN stained cells in the DG. Cells are densely packed at granular layer. Top right, Background-subtracted image by our proposed method. Bottom, Background-subtracted image by Rolling ball algorithm. Note that Rolling ball algorithm subtracted almost all NeuN+ signals because cells are densely distributed within DG. Ball radius parameter used for Rolling ball algorithm is 22.3 µm. The image is in the same section shown in Fig. 4. Scale bars, 50 µm (A) and 200 µm (B).

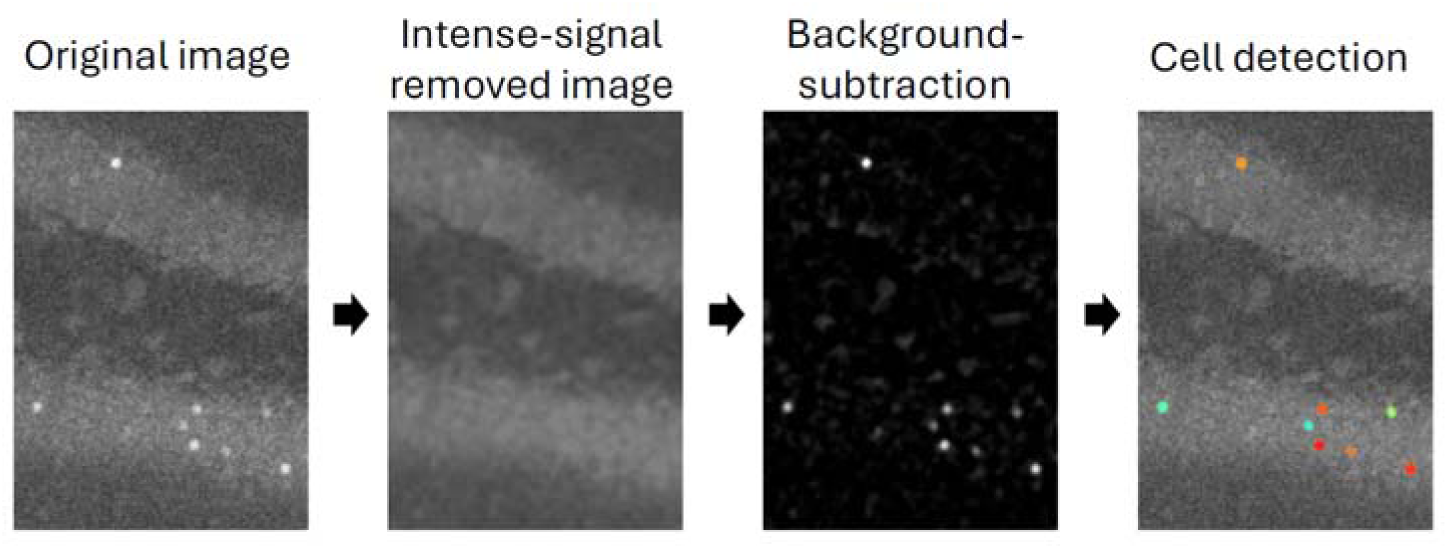
Graphical abstract

